# A four-point measurement probe for brain tissue conductivity and permittivity characterization

**DOI:** 10.1101/2021.04.29.441988

**Authors:** Lucas Poßner, Lydia Seebeck, Matthias Laukner, Florian Wilhelmy, Dirk Lindner, Uwe Pliquett, Bojana Petkovic, Marek Ziolkowski, Thomas R. Knösche, Konstantin Weise

## Abstract

We present a measurement probe consisting of four platinum electrodes fused into soda-lime glass. The probe is based on a four-terminal sensing technique that uses separate pairs of current-carrying and voltage-sensing electrodes and performs more accurate measurements than the simpler and more usual two-terminal sensing technique. The electrodes are aranged within an area of 1 mm^2^ allowing for local characterization of different brain tissues. The operation of the probe is demonstrated in situ on post mortem porcine brain tissue by measuring the impedance spectra of grey and white matter in a frequency range of 1 kHz < *f* < 1 MHz. The conductivity and relative permittivity are derived from the impedance spectra using the geometry factor of the probe. The geometry factor is obtained experimentally by measuring the impedances of an electrolytical dilution series with known conductivities. The obtained conductivity of grey matter is in the order of 0.11 S/m and of white matter is in the order of 0.07 S/m over the acquired frequency range.

## 1. Introduction

The electric properties of in vivo human brain tissue are characterized by its electric conductivity and its relative permittivity. These properties are substantial for the modeling of invasive and non-invasive brain measurement and stimulation techniques, such as EEG, MEG, TMS, TES and DBS [Saturnino et al. 2019, Peterchev et al. 2012, Cho et al. 2015]. Further, these properties can be used for the classification of pathological tissue, such as glioma and other types of brain tumors [Jahnke et al. 2013]. These applications rely on the precise knowledge of the electric material parameters of brain tissue. Previous studies were conducted (i) taking a small number of subjects (≈10); (ii) using inappropriate measurement techniques such as monopolar measurements; (iii) measuring only at one frequency [Latikka et al. 2001, Koessler et al. 2017, Latikka and Eskola 2019]. In a previous study [Poßner et al. 2020] we used a commercial brain stimulation system and a two-point measurement probe to acquire the impedance of post mortem porcine brain tissue and showed that it is possible to distinguish grey and white matter tissue based on the measured impedances. Further investigations have shown that polarization effects at the electrodes of the two-point probe complicate the derivation of conductivity and permittivity values. In this work, we present a proof-of-principle study using a custom-made four-point measurement probe (Kurt-Schwabe-Institut für Meß- und Sensortechnik Meinsberg e.V., 04736 Waldheim, Germany) to obtain the conductivity and relative permittivity spectra of grey and white matter of post mortem porcine brain.

## 2. Methods

### 2.1 Measurement setup

The probe consists of four platinum electrodes fused into soda-lime glass. The probe and an enlarged section of the tip are shown in Fig. 1. The electrodes are used in a four-point configuration with two current-carrying electrodes (E2, E3) and two voltage-sensing electrodes (E1, E4). This configuration minimizes the influence of the electrode polarization. The probe was connected to a commercial impedance analyzer (ISX-3 mini, Sciospec Scientific Instruments GmbH, 04828 Bennewitz, Germany). The impedances were measured within a frequency range of 1 kHz < *f* < 1 MHz and with a current amplitude of *Î* = 100 μA.

**Figure 1.**
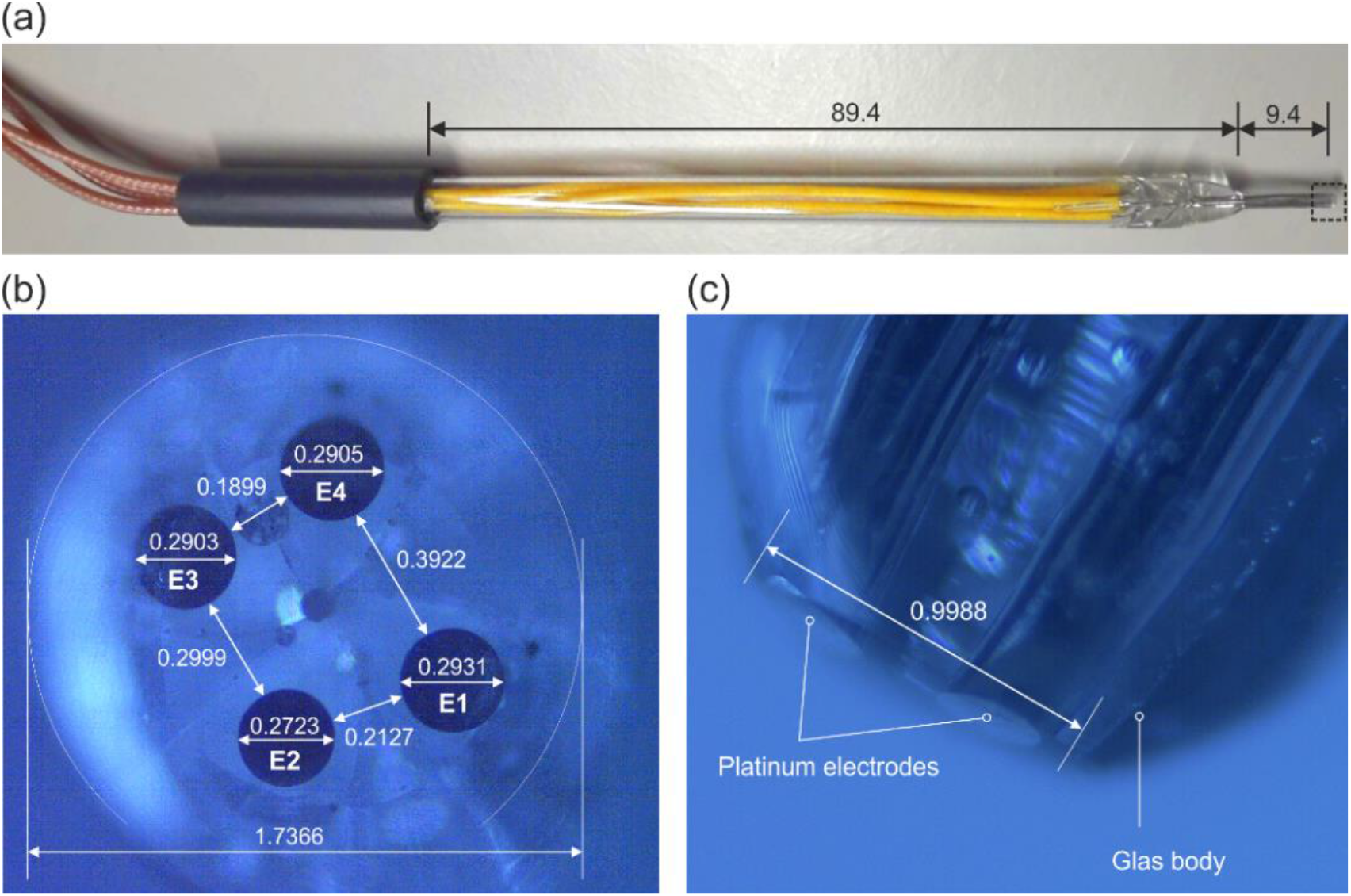
(a) Measurement probe consisting of four platinum electrodes (E1-E4) fused into soda-lime glass; (b) Magnification of the probe tip (frontal view); (c) Magnification of the probe tip (side view). (All measures in mm)

### 2.2 Measurement of post mortem porcine brain impedance

The measurements were performed on post mortem porcine brain at its in situ position inside the head. The pig was regularly slaughtered in a slaughterhouse using electric stunning and the head was separated from the body. By the time of the measurement, it was dead for less than 2 hours. We performed a craniotomy on the porcine head, which left the brain undamaged. The cortex was cut with a scalpel and the impedances of grey and white matter were measured.

### 2.3 Derivation of the conductivity and relative permittivity

The admittance *<u>Y</u>*= 1/*<u>Z</u>* of biological tissue is assumed to not have any inductive properties. Therefore, it can be written as *<u>Y</u>*= *G* + *jωC*, where *G* is the conductance and *C* is the capacitance of the tissue. This can be written as the quotient of the complex conductivity <u>*σ*</u> and a geometry factor *k*: *<u>Y</u>*=<u>*σ*</u>/*k* = (*σ* + *jωε*_0_*ε*_*r*_)/*k*. The geometry factor *k* depends on the geometry of the probe and the electric field distribution in the tissue. Using this relationship, the complex conductivity <u>*σ*</u> can be obtained from the measured admittance *<u>Y</u>*. This procedure was calibrated by obtaining the geometry factor experimentally. An electrolytical dilution series of saline was prepared with an initial concentration of *c* = 194 m mol/l. The solution was diluted 5 times. After each step, the conductivity was measured using a conductivity meter (Cobra SMARTsense, PHYWE Systeme GmbH und Co. KG, 37079 Göttingen, Germany). The concentrations of the solutions and the corresponding conductivities are shown in Table 1.

**Table 1.** Concentration and conductivity of the prepared solutions

| Dilution | Initial | 1 | 2 | 3 | 4 | 5 |
| --- | --- | --- | --- | --- | --- | --- |
| Concentration/(m mol/l) | 194 | 97 | 48.5 | 24.3 | 12.1 | 6.1 |
| Conductivity/(S/m) | 2.071 | 1.041 | 0.521 | 0.266 | 0.136 | 0.073 |

We used the probe to measure the admittance of the solutions within a frequency range of 100 Hz < *f* < 10 kHz. The absolute value of the admittance |*<u>Y</u>*| was constant and linear with respect to the conductivity *σ* and the phase angle *φ* was below 5°. Therefore, the susceptance of the admittance is neglected, and it can be set *<u>Y</u>*= *G*. The geometry factor *k* is calculated by *k* = *σ*/*G*. The obtained values are shown in Fig. 2. The geometry factor *k* is calculated as *k* = 219.78 1/m.

**Figure 2.**
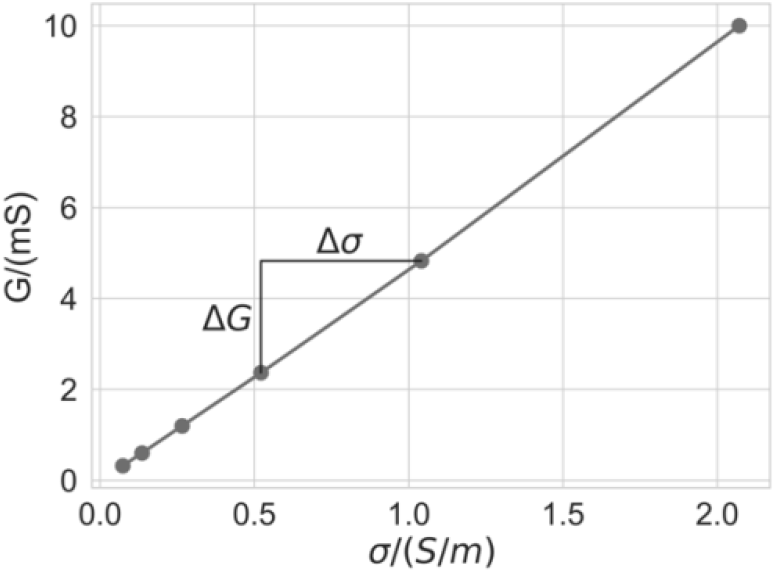
Relationship between the measured conductance and the conductivity of the solutions.

## 3. Results

The impedances of grey and white matter were obtained in a measurement series with ten repetitions. This included the repeated application of the measurement probe on the respective tissues. The absolute values of the impedances and the corresponding phase angles are shown in Fig. 3(a) and (b), respectively, for both grey and white matter. The impedances differ significantly between the tissues (*p* ≪ 0.001, *f* = 10 kHz). The electric conductivity and the relative permittivity of grey and white matter are calculated from the measured impedances by applying the previously obtained geometry factor *k*. The results are shown in Fig. 3(c) and (d), respectively.

**Figure 3.**
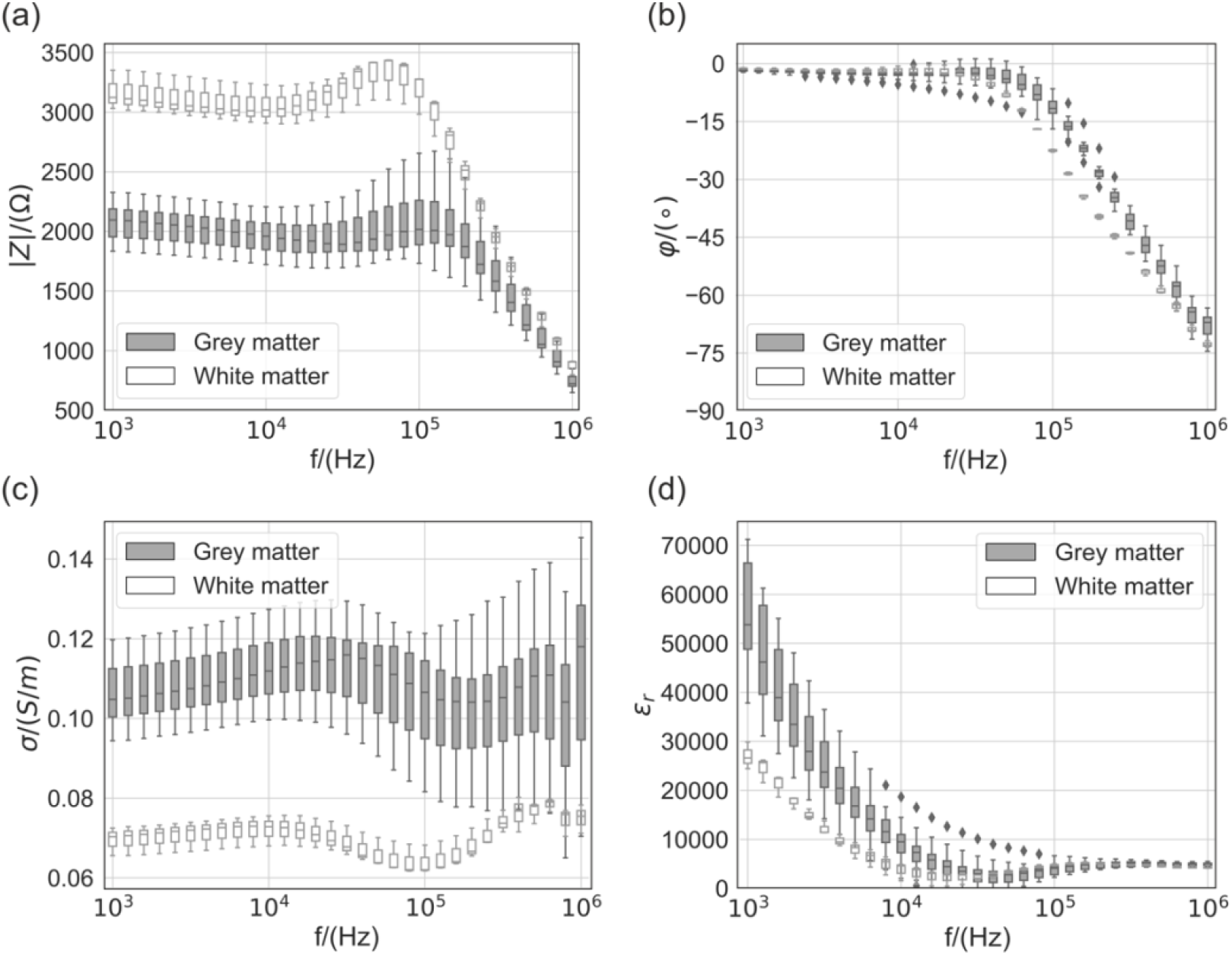
(a) Absolute values of the obtained impedances; (b) Phase angles of the obtained impedances; (c) Calculated conductivities; (d) Calculated permittivities.

## 4. Discussion

The obtained conductivities of grey and white matter are in the order of 0.11 S/m and 0.07 S/m, respectively, over the acquired frequency range. These values are in the range of reported literature values obtained from human brain tissue (IT’IS database, 10 kHz: GM: 0.115 S/m; WM: 0.0695 S/m and Wagner et al. (2004), GM: 0.276 S/m; WM: 0.126 S/m). Differences can be explained by the different species and that our measurements were performed post mortem. The obtained relative permittivity values are lower than the reported literature values (IT’IS database, 10 kHz: GM: 22200; WM: 12500). Further, negative values were obtained at frequencies 10 kHz < *f* < 50 kHz. This is due to the inductive behavior of the measured impedance in this frequency range which could arise from leakage currents through stray capacitances in the measurement chain. It can be observed that the variability between repeated applications of the measurement probe is higher for grey matter. The reason might be the inhomogeneity of this tissue originating from the layered structure of the cerebral cortex. It can be observed that the variability between repeated applications of the measurement probe is higher for grey matter. The reason might be the inhomogeneity of this tissue originating from the layered structure of the cerebral cortex.

## 5. Conclusions

With the presented four-point measurement probe it is possible to perform reliable impedance measurements. To obtain significant conductivity and permittivity values, the acquired impedances need to be compensated with respect to the electrical properties of the measurement chain (measurement fixtures, measurement probe) which is subject of future work.

## Notes

### Competing Interest Statement

The authors have declared no competing interest.

